# A Genetically Encoded Trimethylsilyl 1D ^1^H-NMR Probe for Conformation Change in Large Membrane Protein Complexes

**DOI:** 10.1101/2019.12.18.873729

**Authors:** Qi Liu, Qing-tao He, Xiao-xuan Lyu, Fan Yang, Zhong-liang Zhu, Peng Xiao, Zhao Yang, Feng Zhang, Zhao-ya Yang, Xiao-yan Wang, Peng Sun, Qian-wen Wang, Chang-xiu Qu, Zheng Gong, Jing-Yu Lin, Zhen Xu, Shao-le Song, Shen-ming Huang, Sheng-chao Guo, Ming-jie Han, Kong-kai Zhu, Xin Chen, Alem W. Kahsai, Kun-Hong Xiao, Wei Kong, Xiao Yu, Ke Ruan, Fa-hui Li, Xiao-gang Niu, Chang-wen Jin, Jiangyun Wang, Jin-peng Sun

## Abstract

While one dimensional ^1^H nuclear magnetic resonance (1D ^1^H-NMR) spectroscopy is one of the most important and convenient method for measuring conformation change in biomacromolecules, characterization of protein dynamics in large membrane protein complexes by 1D ^1^H-NMR remains challenging, due to the difficulty of spectra assignment, low signal-to-noise ratio (S/N) and the need for large amount of protein. Here we report the site-specific incorporation of 4-trimethylsilyl phenylalanine (TMSiPhe) into proteins, through genetic code expansion in *Escherichia coli* cells, and the measurement of multiple conformational states in membrane protein complex by 1D ^1^H-NMR. The unique up-field ^1^H-NMR chemical shift of TMSiPhe, highly efficient and specific incorporation of TMSiPhe enabled facile assignment of the TMSiPhe ^1^H-NMR signal, and characterization of multiple conformational state in a 150 kilodalton (kD) membrane protein complex, using only 5 μM of protein and 20 min spectra accumulation time. This highly efficient and convenient methods should be broadly applicable for the investigation of dynamic conformation change of protein complexes.

## Introduction

Membrane proteins account for about 30% of all proteins in living cells, and play critical roles such as material transportation and signal transduction. Because of their critical function in physiological processes, membrane proteins have become one of the most attractive research areas in biochemistry, biophysics and pharmaceutical industry. Indeed, knowledge about the structure and dynamics of membrane proteins is essential for effective drug design^1–3^. However, characterization of multiple functionally important conformation states in large membrane protein complexes has remained to be very challenging^4–6^.

Solution NMR is a powerful tool for studying the structure and dynamics of membrane protein complexes. The developments of isotopic labeling strategies and muti-dimensional NMR methods have facilitated the study of protein complexes up to 1 million Dalton^7^. However, the application of these methods, especially the assignment of complicated multidimensional spectrum is expensive and technically challenging for most biochemistry laboratories. It has recently emerged that one dimensional ^19^F nuclear magnetic resonance (1D ^19^F-NMR) is a powerful and facile method for studying dynamic conformation changes and post-translational modification of proteins, including tyrosine kinases and G-protein coupled receptors (GPCRs), which are among the most important drug targets^8–12^. The advantage of this method is that, through site-specific labelling, typically only one peak is present in the 1D NMR spectra. This allows for the facile characterization of dynamic conformation change with residual precision, without requiring for time-consuming and tedious NMR signal assignment. Despite this significant progress, ^19^F-NMR requires large amount of protein (usually more than 100 μM), and each measurement generally takes more than 12 hours. Therefore, the development of a new chemical biological approach for examination of the conformational dynamics of transmembrane protein complexes using a low concentration of protein is urgently required. However, the large chemical shift anisotropy (CSA) of ^19^F, and the need for expensive ^19^F cryoprobes limit the application of 1D ^19^F-NMR to relatively low-molecular weight proteins. Moreover, this method typically requires more than 50 μM of protein samples and overnight spectra accumulation time. To address these challenges, Otting and colleagues have recently reported one-dimensional ^1^H NMR (1D ^1^H-NMR) tert-butyltyrosine probes. While the nine proton singlet from the tert-butyl group give rise to strong ^1^H-NMR signals, its chemical shift around 1.3 ppm overlaps strongly with the methyl group ^1^H-NMR signals of proteins, and is often difficult to assign. By contrast, the ^1^H-NMR signal from trimethylsilyl (TMS) group has a chemical shift around 0 ppm, that is free of other ^1^H-NMR signal typically present in proteins. Using a cell-free translation system, and a low-efficiency, promiscuous cyanophenylalanine-tRNA synthetase, Otting *et al.* reported the site-specific labelling of proteins using 4-(trimethylsilyl)phenylalanine (TMSiPhe). However, it was observed that ^1^H-NMR signal from TMSiPhe in labelled protein was about ten times smaller than expected, which may be attributed to limited compatibility of the cyanophenylalanine-tRNA synthetase with TMSiPhe. Indeed, no mass spectrometry result was shown to delineate the identity of the genetically incorporated amino acid, in response to UAG codon. Moreover, it was stated that cell-free translation system works best for proteins smaller than 50 kDa, and site-specific labelling of TMSiPhe on larger proteins was unsuccessful, presumably due to the lack of protein chaperons.

To make the 1D ^1^H-NMR method broadly applicable for the investigation of dynamic conformation change for both small proteins and large membrane protein complexes, here we report the highly efficient and selective incorporation of TMSiPhe in proteins in *E. coli* cells. Key for the success is the identification of a mutant *Methanococcus jannaschii* tyrosyl tRNA synthetase (TyrRS), which exhibits high activity and specificity toward 4-trimethylsilyl phenylalanine (TMSiPhe) in *E. coli* cells. Notably, crystallographic analysis revealed structural changes that reshaped the TMSiPhe-specific amino-acyl tRNA synthetase (TMSiPheRS) to accommodate the large trimethylsilyl (TMS) group. Through sodium dodecyl sulfate–polyacrylamide gel electrophoresis (SDS-PAGE) analysis and mass spectrometry (MS), we show that TMSiPhe is genetically incoporated into protein selectively by the UAG codon, and that the UAG codon only encode TMSiPhe. This was not demonstrated previously^13^. Due to the high efficiency and fidelity of TMSiPheRS, we characterized multiple conformational state in a 150 kilodalton (kD) membrane protein complex, using only 5 μM of protein and 20 min spectra accumulation time. We then applied this method to investigate the activation mechanism of arrestin, an important signal transducer downstream of most G-protein-coupled receptors (GPCRs)^12,14–22^.

## Results

### Discovery of a TMSiPhe-specific amino-acyl tRNA synthetase (TMSiPheRS)

The genetic code expansion technique has been widely used recently to incorporate unnatural amino acids at specific positions in a protein to enable in-depth investigation of many important biological processes^23–25^. Such a system includes a synthetic unnatural amino acid (UAA), an orthogonal aminoacyl-tRNA synthetase (aaRS)-tRNA pair derived from directed evolution, and a host protein production organism^23–25^. We synthesized TMSiPhe to facilitate TMS group incorporation into protein, through an optimized route (Figure S2)^26^. TMSiPhe was then used for selection of a tRNA synthetase which accommodate this UAA, using a mutant library of *Methanococcus jannaschii* tyrosyl tRNA synthetase (*Mj*-TyrRS)^27,28^. The mutant library of the *Mj*-TyrRS was designed by randomizing six active site residues (Y32, L65, F108, Q109, D158 and L162) that were within 6.5 Å of the tyrosine substrate, and by performing mutating one of the six residues I63, A67, H70, Y114, I159, and V164 to G, or keeping these residues unchanged as previously described^29^. In the positive selection, cell survival was dependent on the suppression of an amber mutation in the chloramphenicol acetyltransferase gene in the presence of TMSiPhe. By contrast, cells were eliminated if amber codons in the barnase gene was suppressed by natural amino acids in the negative selection without TMSiPhe. Following three rounds of positive selection and two rounds of negative selection^27^, a mutant *M. jannaschii* tyrosyl tRNA synthetase (*Mj*-TyrRS) with specific activity toward TMSiPhe, termed TMSiPheRS, was identified. Sequence analysis revealed that the evolved TMSiPheRS harbors the mutations Tyr32His, Ile63Gly, Leu65Val, His70Gln, Asp158Gly, Ile159Gly and Val164Gly compared to wild-type *Mj*-TyrRS (Figure S3).

We next incorporated TMSiPhe into β-arrestin-1, a signaling protein used here as a model system for evaluation of the TMSiPheRS method. Protein expression was carried out in the presence of β-arrestin-1-H295TAG plasmid, and the pEVOL-TMSiPheRS plasmid (which encodes both TMSiPheRS and *Mj*tRNA_CUA_^Tyr^) in *E. coli* grown in LB media, supplemented with 1 mM TMSiPhe. As negative controls, β-arrestin-1 was also expressed in the absence of any UAA, or in the presence of 1 mM other TMS group containing UAA (TMSiM-dcTyr, TMSiM-Cys, TMSiM-hCys, TMSiM-Tyr). As shown in Figure 1, full-length β-arrestin-1-H295TMSiPhe was expressed in good yield (around 1 mg/L after Ni-NTA affinity column purification), but no full length protein was expressed in the absence of UAA, or in the presence of other TMS group containing UAA, suggesting that TMSiPheRS exhibited good selectivity and activity for TMSiPhe (Figure 1b). Mass spectrometric analysis unambiguously showed the incorporation of TMSiPhe at H295 position in β-arrestin-1 with 100% selectivity (Figure 1c, Figure S4 and Table S1). To further demonstrate the efficiency and selectivity of TMSiPheRS, we expressed green fluorescent protein (GFP) harboring TMSiPhe in the Y182 positon (GFP-Y182TMSiPhe). Crystallization of GFP-Y182TMSiPhe takes approximately 1 week at 16 ℃, followed by X-ray diffraction in Shanghai synchrotron facility. As Figure 1d shows, the TMS group electron density is clearly resolved in the GFP-Y182TMSiPhe crystal structure. These results further demonstrate the high efficiency and fidelity of TMSiPheRS mediated TMSiPhe incorporation, and that TMSiPhe exhibit good stability and compatibility with proteins, which are important for protein structure studies. We then inspected the 1D ^1^H-NMR properties of the TMSiPhe decorated protein. 1D ^1^H-NMR NMR spectroscopy of β-arrestin-1-H295TMSiPhe revealed a unique ^1^H-NMR peak at 0.25 ppm, which is well separated from the other endogenous ^1^H-NMR signals from β-arrestin-1, providing a distinct NMR probe for the examination of the structural dynamics of a specific site (Figure 1e and Figure S5). Notably, the ^1^H-NMR signal of TMSiPhe-incorporated arrestin can be detected at concentrations as low as 5 μM very rapidly (in less than 20 min) using a 950 MHz NMR spectrometer. This high sensitivity is due to the nine equivalent proton present in the TMS group, and the usage of a high-field NMR spectrometer. By contrast, the large chemical shift anisotropy (CSA) of ^19^F in general limit the ^19^F-NMR studies of proteins to NMR spectrometers lower than 600 MHz. The high sensitivity of the TMSiPhe probe is important for 1D ^1^H-NMR studies, especially for large membrane proteins complexes since they are prone for aggregation in high concentration.

**Figure. 1.**
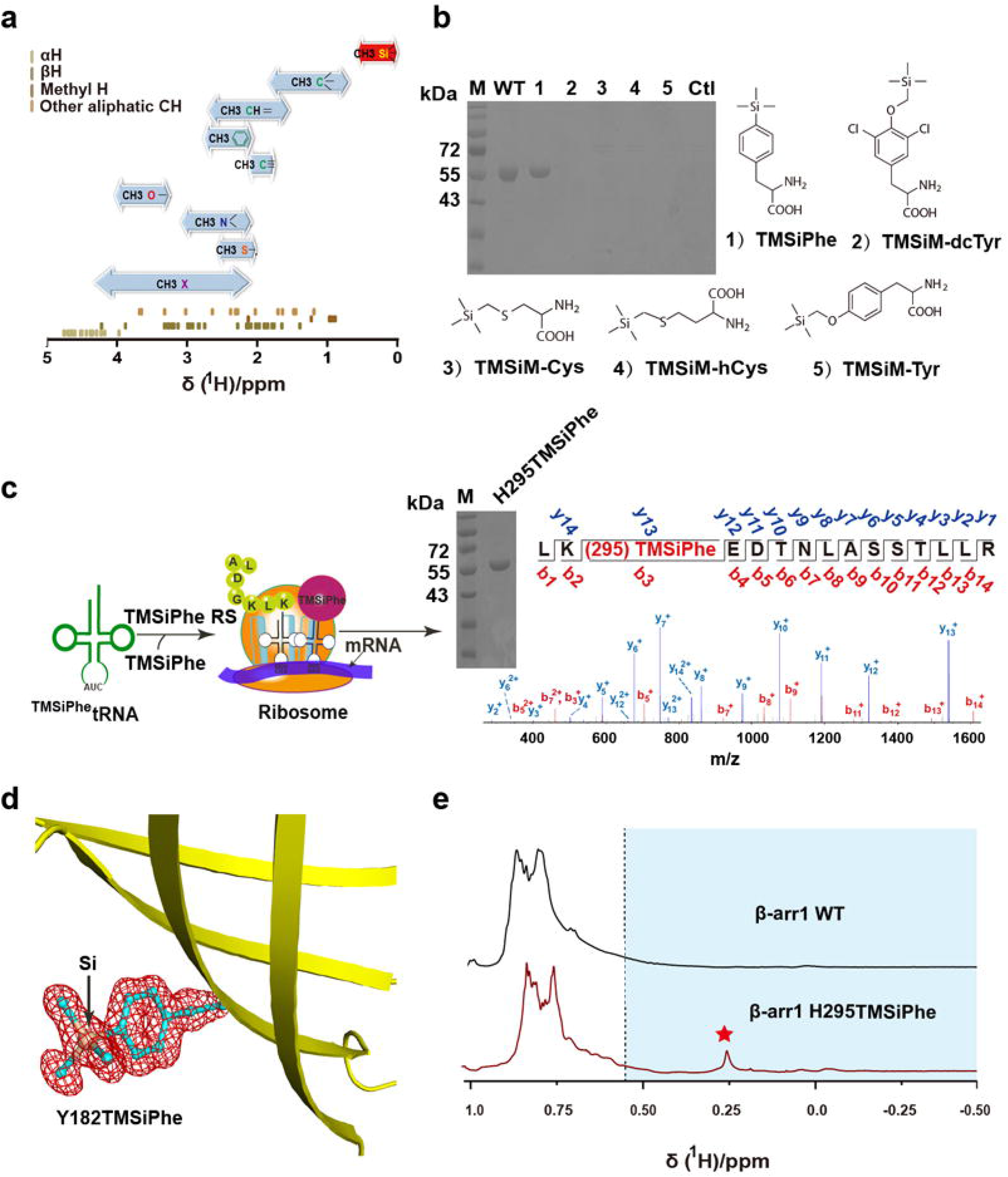
Development of TMSiPheRS by genetic code expansion and the selectivity of TMSiPheRS. **(a)** The ranges of the methyl ^1^H chemical shifts^53^(shown by bidirectional arrows) and the distribution of random-coil aliphatic CH ^1^H chemical shifts for the 20 genetically coded amino acids^54^. The ^1^H chemical shifts of methyl silicon group are specified in red. **(b)** Coomassie-stained gel analysis of full-length β-arrestin-1 expression in *E. coli* cells that were cotransfected with the β-arrestin-1-H295TAG plasmid and the pEVOL-TMSiPheRS plasmid, encoding a specific *M. jannaschii* tyrosyl amber suppressor tRNA/tyrosyl-tRNA synthetase mutant grown in the presence or absence of different silicon-containing compounds. WT indicates wild-type arrestin without any change in genetic code. Full-length β-arrestin-1 protein was obtained only in the presence of TMSiPhe for TAG mutation of β-arrestin-1 or WT. These results suggested that the evolved TMSiPheRS exhibited significant structural selectivity for TMSiPhe over other silicon-containing chemicals. The chemical abbreviations are as follows:

(1) 4-(trimethylsilyl) phenylalanine, TMSiPhe;
(2) 3,5-dichloro-4-[(trimethylsilyl) methoxy]phenylalanine, TMSiM-dcTy;
(3) 2-amino-3-((trimethylsilyl)methylthio)propanoic acid, TMSiM-Cys;
(4) 2-amino-4-((trimethylsilyl)methylthio)butanoic acid, TMSiM-hCys;
(5) 4-[(trimethylsilyl)ethoxy]phenylalanine, TMSiM-Tyr;
(6) control, Ctl. There were no unnatural amino acids added to the culture. **(c)** Schematic flowchart for the incorporation of TMSiPhe into β-arrestin-1 at the H295 site. Full-length β-arrestin-1 protein was obtained by cotransfection with the β-arrestin-1 H295 TAG mutant plasmid and the pEVOL-TMSiPheRS plasmid, with TMSiPhe supplementation of the culture medium. The purity of the protein was determined by electrophoresis. The protein was subjected to trypsin digestion and analyzed by MS/MS. These results unambiguously confirmed that TMSiPhe was selectively incorporated into β-arrestin-1 at the H295 position. m/z, mass/charge ratio. **(d)** The 2Fo-Fc annealing omit map of sfGFP-Y182-TMSiPhe clearly shows the electron density of TMSiPhe. The map was contoured at 1.1 σ. **(e)** 1D ^1^H NMR spectra for the β-arrestin-1 H295 TMSiPhe mutant were compared with those for wild-type β-arrestin-1 cultured in the presence of TMSiPhe. The spectra were recorded in a buffer containing 50 mM Tris-HCl (pH 7.5) and 150 mM NaCl at 25°C using a Bruker 950 MHz NMR spectrometer. The β-arrestin-1 H295 TMSiPhe chemical shift at 0.26 ppm was consistent with the predicted chemical shift of the TMSi group. The ^1^H NMR signals of TMS group substituted amino acids in a protein were generally located in the high-field region (<0.55 ppm, blue area).

### Molecular basis of the selective recognition of TMSiPhe by TMSiPheRS

To investigate the molecular basis of the selective recognition of TMSiPhe by TMSiPheRS, we crystallized TMSiPheRS and analyzed the structures by X-ray crystallography. The crystal structures of TMSiPheRS alone and the complex of TMSiPheRS with TMSiPhe were determined at 1.8 Å and 2.1 Å, respectively (Table S2). The 2Fo-Fc annealing omit map of the TMSiPheRS/TMSiPhe complex unambiguously assigned the electron density for TMSiPhe (Figure 2a). Introduction of the TMS group significantly increased the volume of the amino acid substrate by approximately 60% (Figure 2b). To compensate for this substantial change in volume, three residues, namely, Asp158, Ile159 and Val164, were replaced by the smallest amino acid Gly, and Tyr32 and Leu65 were substituted by the relatively small residues His32 and Val65, respectively (Figure 2b-2c and Figure S3). We then compared the crystal structure of the TMSiPheRS/TMSiPhe complex with that of apo TMSiPheRS. Compared to the structure of TMSiPheRS alone, we observed a dramatic 120-degree rotation of histidine 32 in response to TMSiPhe binding. Moreover, Leu162 rotated approximately 42 degrees to form hydrophobic interactions with the methyl groups of TMSiPhe (Figure 2d). Altogether, Gly34, Val65, Gln70, Phe108, Gln109, Tyr151, Gln155, Gly158, Gly159, Gln173 and His177 defined a hydrophobic pocket for the accommodation of and specific interactions with the phenyl ring and TMS group of TMSiPhe (Figure 2c and Figure S3). These observations provided a structural basis for specific and efficient incorporation of TMSiPhe using evolved TMSiPheRS.

**Figure. 2.**
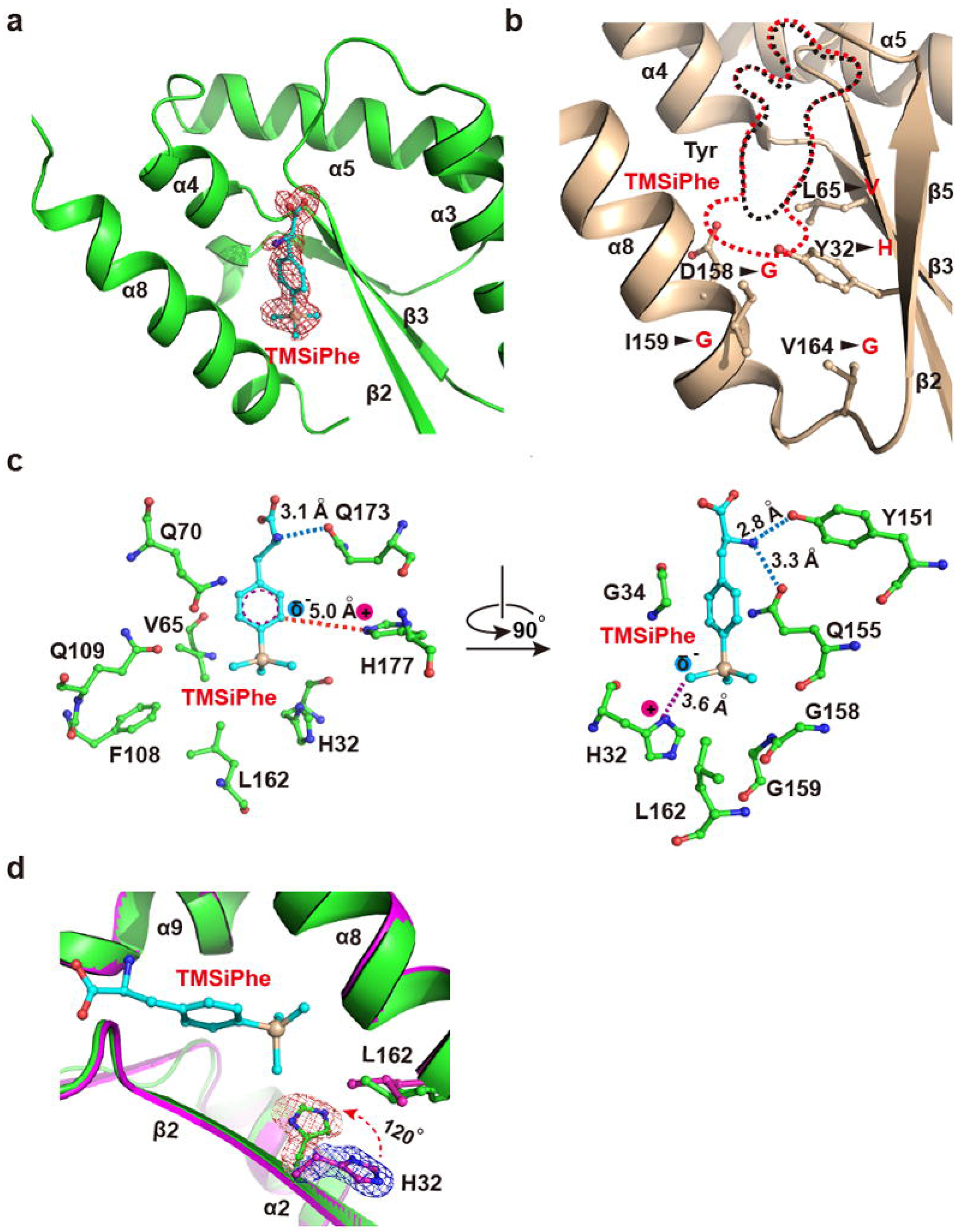
Structural basis for the selective recognition of TMSiPhe by TMSiPheRS. **(a)** Binding of TMSiPhe at the active site of TMSiPheRS. The 2Fo-Fc annealing omit electron density map of TMSiPhe was contoured at 1.0 σ. **(b)** Comparison of the unnatural amino acid-binding pockets between TMSiPheRS (red dotted line) and the wild-type *Mj*-TyrRS (black dotted line, PDB:1J1U). Five key mutations, indicated by arrows, increased the size of the TMSiPhe-binding pocket substantially. **(c)** Interactions between TMSiPhe and TMSiPheRS. The specific interactions include hydrogen bonds (blue dotted line), π-cation interactions (red dotted line), ion-dipole interactions (magenta dotted line) and hydrophobic interactions with surrounding residues (left panel). **(d)** The H32 residue in the β2 strand was rotated approximately 120 degrees in the TMSiPhe/TMSiPheRS complex (green) compared with TMSiPheRS alone (magenta), leading to favorable charged interactions with TMSiPhe.

### Incorporation of TMSiPhe at specific sites in β-arrestin-1 enabled characterization of β-arrestin-1 activation

We then incorporated TMSiPhe into functionally relevant structural motifs of β-arrestin-1, the key signal transducer downstream of almost all 800 GPCRs encoded in the human genome, which functions not only by desensitizing membrane receptors but also by mediating independent downstream signaling after receptor activation^12,14–22,30^(Figure 3a and 3b). Although the functions of many arrestin-mediated receptors have been identified and certain motifs of arrestin are suspected to be involved in specific signaling pathways (Table S3), the correlation between the conformational states of these arrestin motifs and selective receptor functions remains to be elucidated. Incorporation of TMSiPhe into β-arrestin-1 at specific positions, including the receptor-phosphate-binding site (Y21), the finger loop (Y63), the hinge region (Y173), the β-strand XVI (Y249), the loop between β-strands XVIII and XVIIII (R285), the lariat loop (H295) and the C-terminal swapping region (F388), led to unambiguous assignment of NMR peaks between −0.3 ppm and 0.3 ppm in the ^1^H-NMR spectrum (Figure 3b, Figure S6-S7 and Table S4). These positions were proposed to be associated with specific arrestin functions, including receptor or IP6 interactions, or the activation of downstream ERK or AP2 but have never been fully characterized by biophysical methods (Table S3). Therefore, TMSiPhe-containing β-arrestin-1 proteins provide a useful tool for monitoring conformational changes in arrestin in response to receptor activation or other stimuli. For example, the ^1^H-NMR spectrum of native β-arrestin-1 F388-TMSiPhe exhibits a peak at −0.05 ppm, which can be easily identified. By contrast, when another UAA, O-tert-butyltyrosine^13,31^, was genetically encoded into the same position, the peaks for which cannot be assigned due to strong overlap with the methyl signals from the protein (Figure S8). Notably, in response to stimulation with increased concentrations of phospho-vasopressin-2 receptor-C-tail peptide (V2Rpp), the peak at −0.05 ppm gradually disappears, whereas a ^1^H-NMR peak at 0.15 ppm appears, reflecting the transition of the “inactive” arrestin conformation to an “active” arrestin conformation at the F388 position, through dislodgement of this specific C-terminal swapping segment (Figure 3c and 3d). Moreover, the Scatchard plot for the titration experiment performed to examine the binding of V2R-phospho-C-tail to β-arrestin-1 exhibits a straight line with a regression coefficient of 0.99. The calculated *K*_D_ value for the interaction of V2Rpp with β-arrestin-1 was 6.9 ±0.2 μM (Figure 3e and Figure S9). Here, we demonstrate that while the genetic incorporation of TMSiPhe introduce little perturbation to the target protein, it can be used as a convenient tool for determining protein/peptide binding affinities. Since the 1D ^1^H-NMR spectra contain only two peaks which represent the “active” and “inactive” conformation, it takes no effort to perform NMR spectra assignment. Moreover, since ^1^H-NMR is easily accessible to most universities, our method is broadly applicable to most biochemistry laboratories, without requiring for a strong expertise in NMR spectra assignment and multidimensional NMR experiments.

**Figure. 3.**
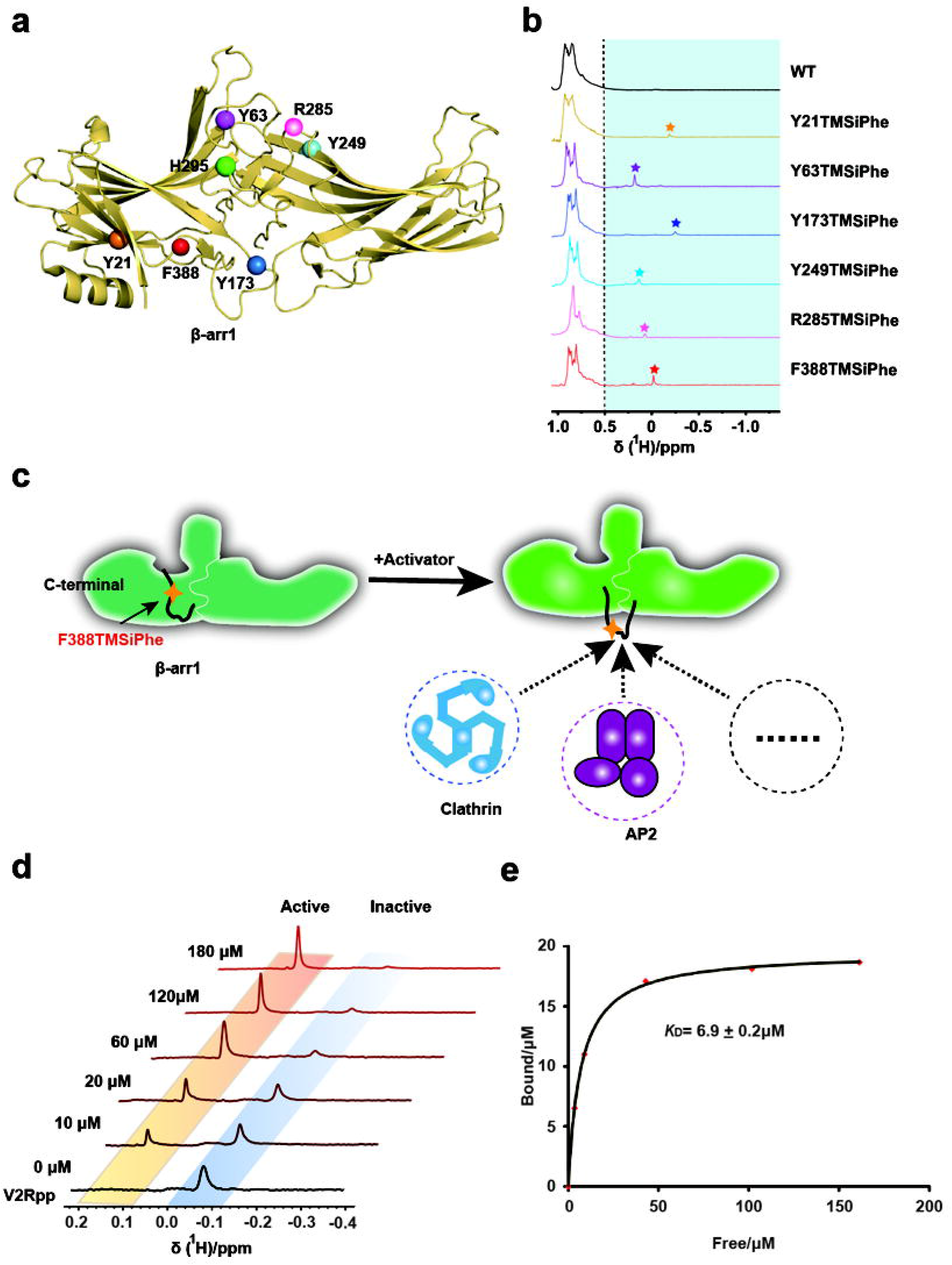
Incorporation of TMSiPhe at different functionally relevant motifs of β-arrestin-1 and characterization of β-arrestin-1 activation by TMSiPheRS. **(a)** Frontal view of the TMSiPhe incorporation sites depicted by spheres in the inactive β-arrestin-1 crystal structure (PDB: 1G4M). Orange, Y21 in the three elements; purple, Y63 in the finger loop; blue, Y173 in the hinge region; cyan, Y249 in β-strand XVI; pink, R285, green, H295 in the lariat loop; red, F388 in the C-terminal swapping segment. **(b)** 1D ^1^H NMR spectra of β-arrestin-1 labeled as described in (3a). The spectra were recorded in a buffer containing 50 mM Tris-HCl (pH 7.5 and 150 mM NaCl at 25°C using a Bruker 950 MHz NMR spectrometer. The protein concentrations were 5~15 μM, and the total recording time per spectrum was 6~15 min. The chemical shift for the TMSiPhe protein was less than 0.55 ppm. **(c)** Cartoon illustration of the activation of β-arrestin-1 and movement of the C-terminal swapping segment of β-arrestin-1. In response to the binding of an activator, such as the phospho-vasopressin receptor C-tail (V2Rpp), the originally embedded C-terminal swapping segment of β-arrestin-1 became highly solvent exposed, thus favoring binding to downstream signaling proteins, for example, clathrin or AP2 (adaptor protein 2). This conformational transition could be monitored by incorporation of TMSiPhe at the F388 position of β-arrestin-1, which is located in the C-terminal swapping segment. **(d)** 1D ^1^H-NMR spectra of β-arrestin-1–F388-TMSiPhe in response to titration with V2Rpp. Two distinct peaks were observed. The peak (−0.05 ppm) representing the inactive state gradually decreased in intensity, while the peak representing the active state (0.15 ppm) steadily increased in intensity. The spectra were recorded in a buffer containing 50 mM Tris-HCl (pH 7.5) and 150 mM NaCl at 25°C using a Bruker 950 MHz NMR spectrometer. **(e)** Analysis of the titration experiments monitored by 1D ^1^H-NMR spectroscopy of β-arrestin-1 F388TMSiPhe (3d). The curve was fitted to the nonlinear regression equation y=B_max_[X]/(*K*_D_+[X], according to the scatchard plot analysis (fig. S9). The *K*_D_ value was calculated at 6.9±0.2 μM (R^2^=0.99).

### Observation of the conformational change of the polar core of β-arrestin-1 by different GPCR ligands through TMSiPhe

Arrestin is known to be activated via both receptor-phosphorylation and active seven transmembrane 7TM core^8,12,15–18,32–37^(Figure 4a). While the recent rhodopsin/visual arrestin complex structure has provided a model of the interactions of the receptor core with arrestin at an atomic resolution^37^, there is little structural information regarding receptor core-induced structural rearrangement of arrestin at the residue level due to technological difficulties in distinguishing the contributions of the receptor core, the receptor-phospho-tail, or the linker and arrestin mutant used in the crystal structures individually, as well as the large amount of the receptor complex required for structural delineation.

**Figure. 4.**
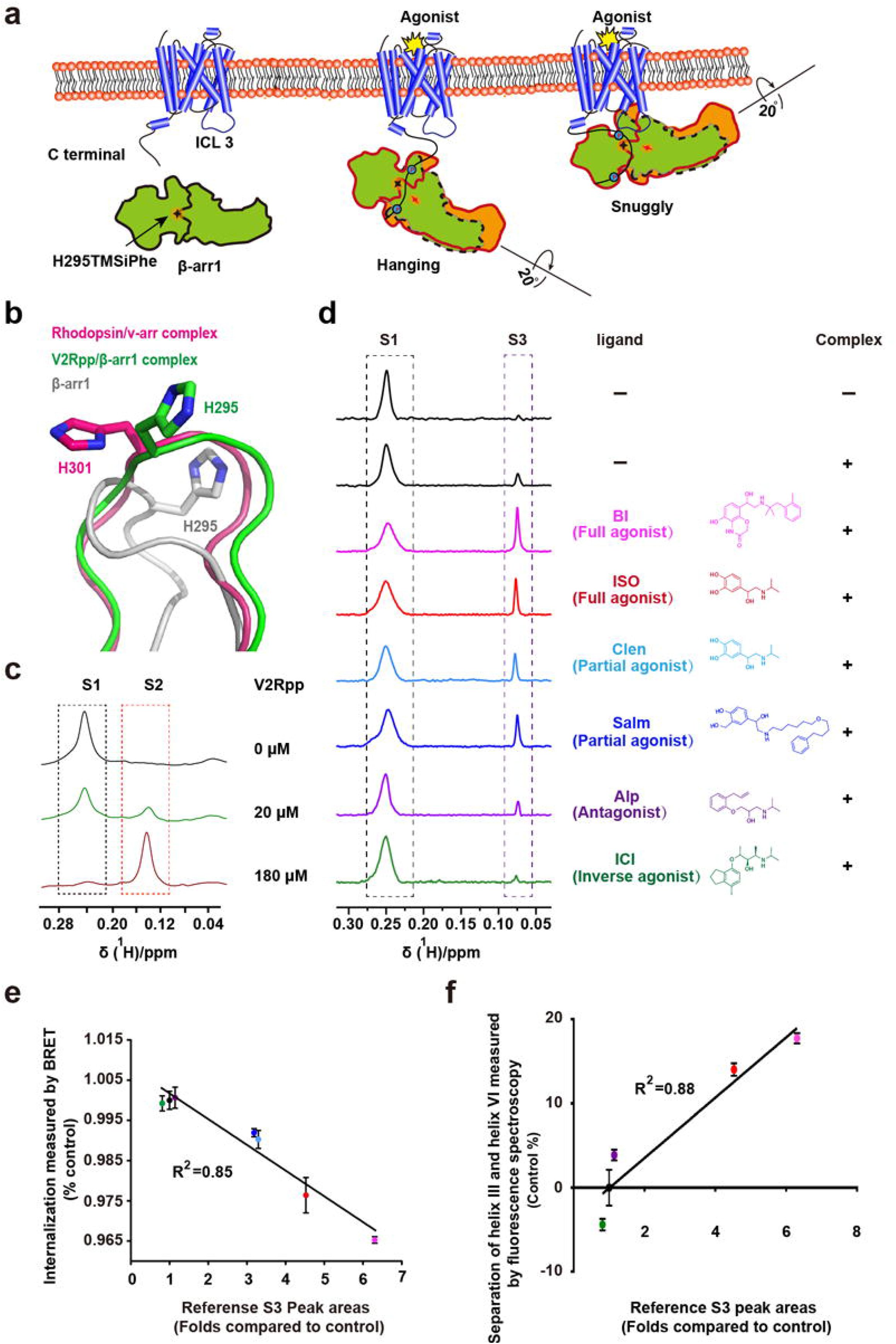
Regulation of the conformational changes of the β-arrestin-1 polar core by different β2AR ligands. **(a)** Cartoon illustration of two distinct interaction modes between GPCRs and β-arrestin-1 (hanging mode and snug mode). The phosphorylated C-tail and transmembrane core of the receptor were each able to independently stimulate arrestin activation. After activation, the N- and C-domains of β-arrestin-1 underwent approximately 20 degrees of rotation. The polar core of β-arrestin-1 is hypothesized to be a critical stabilizer of the inactive state, and introduction of the probe at H295, which is close to this key region, enables detection of conformational changes in the polar core in response to the GPCR activation. **(b)**. Structural comparison of the H295 position in inactive β-arrestin-1 (PDB: 1G4M), the V2Rpp/ β-arrestin-1 complex (PDB: 4JQI) and the rhodopsin/arrestin complex (PDB: 5W0P). The inactive β-arrestin-1 structure is depicted in gray; the V2Rpp/β-arrestin-1 complex is in green; and the rhodopsin-arrestin complex is in red. The two active arrestins have similar conformations at the H295 position, differing significantly from the pose in the inactive arrestin structure. **(c)** 1D ^1^H NMR spectra of β-arrestin-1–H295-TMSiPhe in response to titration with V2Rpp. Two distinct peaks were observed. With increasing concentrations of V2Rpp, the peak at 0.25 ppm decreased (representing the S1 state), whereas a new growing peak was observed at 0.15 ppm (representing the S2 state). **(d)** 1D ^1^H NMR spectra of β-arrestin-1 H295TMSiPhe alone or the ppβ2V2R/β-arrestin-1 H295TMSiPhe/Fab30 complex with or without different ligands and the chemical structures of the ligands used in the current study. After incubation with the phospho-β2AR-V2-tail (ppβ2V2R) and formation of the receptor-arrestin complex, a new NMR signal appeared at 0.07 ppm (designated S3), and the intensity of the S1 peak decreased. When incubated with different β2AR ligands before formation of the ppβ2V2R-β-arrestin-1/Fab30 complex, the S3 state signal intensity of the complex was positively correlated with effects of the ligands on the activation of downstream effectors, such as arrestin. BI, BI-167107; ISO, Isopreteronol; Clen, Clenbuterol; Salm, Salmeterol; Alp, Alprenolol; ICI, ICI-118551. The buffer used for the experiment contained 20 mM HEPES, 150 mM NaCl, 0.01% LMNG, 0.002% CHS, and 10% D2O (pH 7.5 at 25°C). **(e)** Best-fit linear correlation of the peak area representing the amount of the S3 state in the presence of different ligands, with the ligand efficacy for receptor internalization from the BRET experiment in vivo. See fig. S17 for details. **(f)** Best-fit linear correlation of the peak area representing the amount of the S3 state in the presence of different ligands, with the ligand efficacy for separation of the receptor transmembrane III and VI from the TRIQ experiment in vitro^52^. See fig. S18 for details.

Thus, incorporation of TMSiPhe at specific positions in arrestin might facilitate detection of ligand-induced conformational changes in arrestin using 1D ^1^H-NMR. One hallmark of arrestin activation is the approximately 20− degree twist between the N- and C-domains of the protein (Figure 4a). In the inactive state, the N- and C-domains of β-arrestin-1 are tethered by the polar core, which is composed of the extensive charged interactions of Asp26, Arg169, Asp290, Asp297 and Arg393 (Figure S10). Disruption of the salt bridge between Asp297 and Arg393 and that between Asp304 and Arg382, as well as the equivalent rhodopsin-visual arrestin interactions between Asp296 and Arg175 and Asp303 and Arg382, are known to activate arrestin^37^(Figure S10). Notably, the results of recent molecular dynamics studies have indicated that the rotation of Asp296 of visual arrestin (Asp290 in β-arrestin-1) is closely associated with interdomain twisting. We therefore incorporated TMSiPhe at the H295 position, which is close to both D290 and D297 of β-arrestin-1, to monitor the receptor-induced conformational changes in the polar core (Figure 4a-4b and Figure S10). Specific incorporation of TMSiPhe at the H295 position in β-arrestin-1 did not impair the structural integrity of the protein, as H295-TMSiPhe-β-arrestin-1 exhibited normal activation in response to the V2-receptor-phospho-tail interaction (Figure S11). For structural validation of the TMSiPheRS study with H295-TMSiPhe-β-arrestin-1, we performed ^1^H-NMR measurements using the conditions for the crystal structures of β-arrestin-1 in both the inactive apo-arrestin and in active arrestin stabilized by vasopressin 2 receptor phospho-tail (V2Rpp) and the conformationally selective antibody Fab30^35^. Notably, in the “two-step arrestin recruitment model of the receptor”, V2Rpp/β-arrestin-1 mostly exhibited the “hanging” mode, whereas the phospho-receptor/β-arrestin-1 complex encompassing the core interaction represented the “snuggly” mode^32,35^(Figure 4a). Superimposition of the inactive and active arrestin structures revealed that both the “hanging” and “snuggly” modes of active arrestin had similar conformations at the H295 position, differing significantly from the modes of inactive arrestin, which featured considerable movement of the lariat loop (Figure 4b).

In the inactive state, the 1D 1H-NMR spectrum of H295-TMSiPhe-β-arrestin-1 contained mainly one peak at 0.25 ppm, which was designated S1 (Figure 4c). Upon the addition of increasing the concentration of V2Rpp, the peak volume of S1 gradually decreased, accompanied by the growth of a new peak at 0.15 ppm, which was designated S2. The 1D ^1^H-NMR spectrum obtained with a saturating concentration of V2Rpp mainly exhibited an S2 peak, indicating that S2 represented an active state of H295TMSiPhe, whereas S1 represented the inactive state of β-arrestin-1 H295TMSiPhe (Figure 4b and 4c).

We then inspected the conformational change at the H295 site in response to occupation of the receptor by a panel of ligands with the same phospho-receptor-tail by using the β2 adrenergic receptor (β2AR) as a prototypic model. As previously described, we obtained the phospho-β2AR-V2-tail chimera (ppβ2V2R) by stimulating Sf9 cells with ISO (Isoproterenol), a low-affinity ligand, before harvesting the cells and washing out the residual ligands by affinity chromatography^38^. The purified ppβ2V2R was then incubated with various ligands and then used to form a stable receptor/arrestin complex by further incubation with β-arrestin-1 and the conformationally selective antibody fragment Fab30 (Figure S12). Complex formation was verified by size-exclusion chromatography, and 1D ^1^H-NMR was performed to monitor changes in the NMR signal (Figure 4d). Application of the arrestin active conformation stabilizing Fab30 alone had no significant effect on the NMR spectrum of β-arrestin-1 H295TMSiPhe (Figure S13). Notably, upon incubation with ppβ2V2R and Fab30, a new NMR signal appeared at 0.07 ppm (designated S3), which was associated with the decrease in the S1 peak (Figure 4d and Figure S14-S15). Therefore, S3 may represent the active arrestin state of H295TMSiPhe in the presence of ppβ2V2R,. The sharp S3 peak compared to S1 might indicate a highly solvent-exposed structure of the H295 state in β-arrestin-1 after forming the complex with the ppβ2V2R, as observed in the crystal structure of the rhodopsin/visual arrestin complex. ^1^H-NMR chemical shift is sensitive to the change of hydrogen bonding, local dielectric constant, and nearby aromatic residues. Thus, NMR chemical shifts are sensitive to subtle structural changes in proteins. The S2 and S3 states have similar loop structures, but subtle differences in sidechain orientation are obvious (Figure 4b).

We next examined the 1D ^1^H-NMR spectrum of the ppβ2V2R/β-arrestin-1 complex in the presence of various β2AR ligands exhibiting different pharmacological activities. Importantly, while the S1 state population of H295TMSiPhe decreased upon addition of various agonists, including the full agonists ISO and BI-167107, or the partial agonists Clenbuterol (Clen) and Salmaterol (Salm), the S3 state population increased (Figure 4e and Figure S15-S16). By contrast, the neutral antagonist Alprenolol (Alp) showed no effect on the S3 state, whereas the inverse antagonist ICI-118551 (ICI) reduced the population of the S3 state (Figure 4d and Figure S15). Moreover, the volume of the S3 state corresponds to the potency of the ligand in inducing receptor internalization (Figure 4e and Figure S17). These trends mirrored the ability of the ligand to promote torsion between helix VI and helix III, a hallmark of the conformational changes in the receptor 7-transmembrane core induced by agonists (Figure 4f and Figure S18). Overall, the 1D 1H-NMR spectrum of β-arrestin-1 H295TMSiPhe indicated that the conformational changes in the polar core of the arrestin in response to ligand properties are associated with the abilities of these ligands to promote receptor internalization in the presence of the same receptor (Figure 4e-4f and Figure S17).

### The role of the receptor 7TM core in mediating the ligand-regulated conformational change in the polar core of β-arrestin-1

To confirm that the observed S3 signal in the 1D 1H-NMR spectrum was dependent on the interaction of β-arrestin-1 with the receptor core, we performed a competition assay using a well-characterized binding partner of the receptor 7-transmembrane core, namely, the Gα protein C-tail (Gα-CT) (Figure 5a). Moreover, direct engagement of the β-arrestin-1 finger loop and Gα-CT of G-protein serves as a major interaction interface with the receptor 7-transmembrane core, which was supported by recent cross-linking and electron microscopic studies^35,39–41^. Therefore, we prepared the ISO/ppβ2V2R/β-arrestin-1/Fab30 complex in the presence of Gα-CT. Incubation with Gα-CT did not disrupt the ISO/ppβ2V2R/β-arrestin-1/Fab30 complex, as the presence of Gα-CT did not alter the SEC (size-exclusion chromatography) profile (Figure 5b). Notably, while incubation of Gα-CT with β-arrestin-1-H295-TMSiPhe led to no significant alteration in the 1D ^1^H-NMR spectrum, addition of Gα-CT with the ISO/ppβ2V2R/β-arrestin-1/Fab30 complex significantly decreased the S3 state and increased the S1 state, suggesting that the observed S1 state reduction was mainly due to elimination of the receptor core interaction with β-arrestin-1 by the binding of Gα-CT (Figure 5c). Because H295 is located in the close proximity to the polar core residues Asp290 and Asp297 of β-arrestin-1, the 1D ^1^H-NMR spectrum obtained from Gα-CT competition experiments with H295-TMSiPhe-β-arrestin-1 confirmed that the agonist ISO was able to induce conformational changes in the polar core of β-arrestin-1 via direct transmembrane core interactions.

**Figure. 5.**
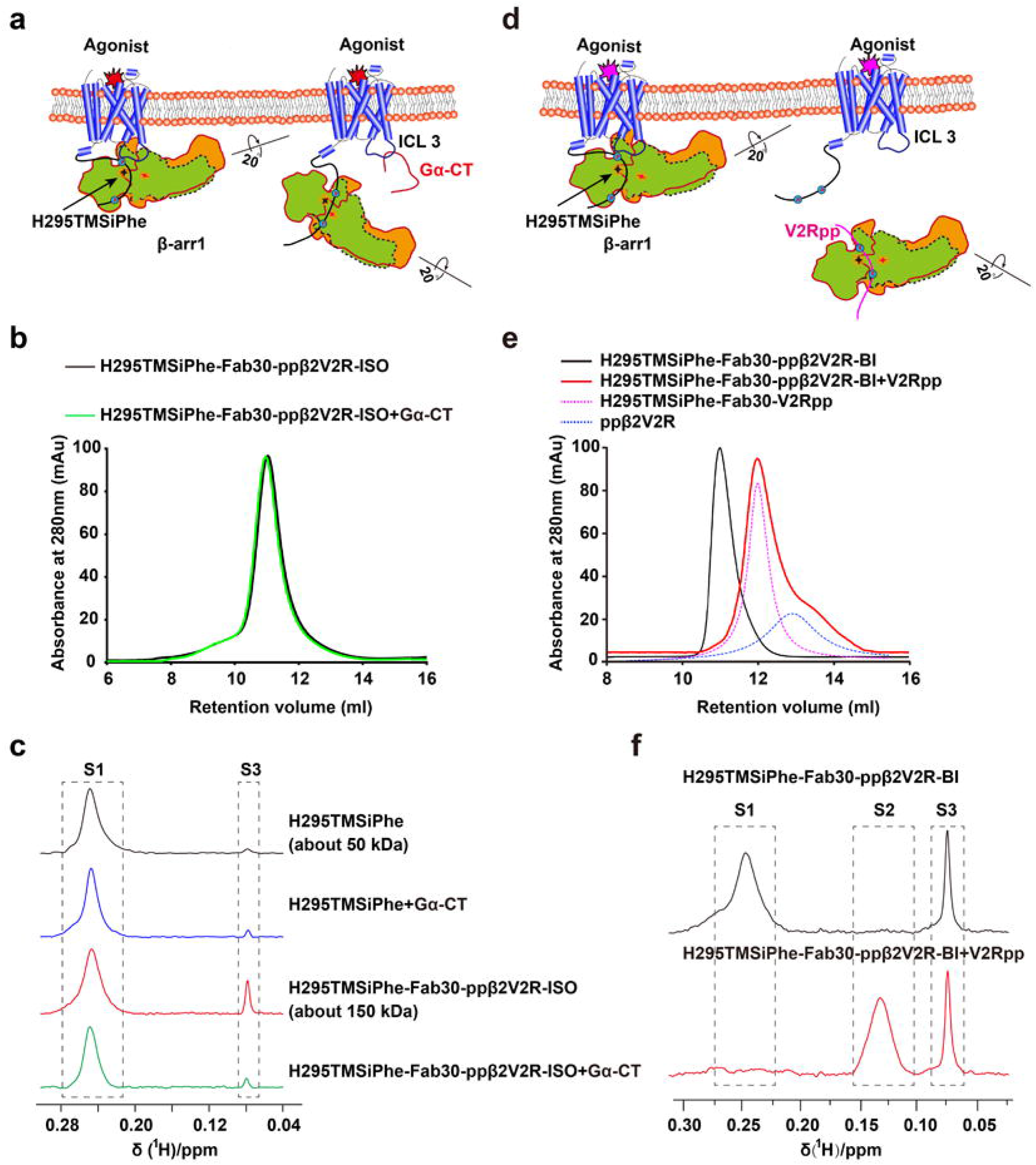
Gα-CT competition experiments confirmed the essential role of the 7TM core of the receptor in mediating the ligand-regulated conformational change in β-arrestin-1. **(a)** Schematic diagram of the interaction between GPCRs and β-arrestin-1 in the presence of excess Gα C-terminus (Gα-CT), which has been described in previous reports^55^. The interaction between β-arrestin-1 and the GPCR TM core was abolished via steric hindrance by Gα-CT. β-Arrestin-1 still interacts with the phosphorylated GPCR C-terminal tail and thus forms a complex with the receptor. **(b)** ISO/ppβ2V2R/β-arrestin-1/Fab30 complex formation was not disrupted by Gα-CT in a size-exclusion assay. The similar SEC profile observed with or without Gα-CT suggests that Gα-CT did not disrupt the ISO/ppβ2V2R/β-arrestin-1/Fab30 complex. Size-exclusion chromatography experiments were performed on an AKTA Purifier equipped with a Superdex 200 (10/300GL) column. Black: ISO/ppβ2V2R/β-arrestin-1-H295TMSiPhe/Fab30 complex, green: ISO/ppβ2V2R/β-arrestin-1-H295TMSiPhe/Fab30 complex mixed with the 200 μM Gα-CT. **(c)** 1D ^1^H NMR spectra of β-arrestin-1-H295TMSiPhe in the presence of Gα-CT. The transformation from S1 to S3 induced by the ISO/ppβ2V2R/β-arrestin-1/Fab30 complex was significantly weakened by the addition of Gα-CT, suggesting the observed S3 state reduction was mainly due to the elimination of the receptor core interaction with β-arrestin-1 by the binding of Gα-CT. **(d)** Schematic diagram of the competing experiments of the ppβ2V2R/β-arrestin-1 complex disrupted by the presence of excess V2Rpp. β-Arrestin-1 dissociated from the phosphorylated β2V2R due to competition with V2Rpp. **(e)** ISO/ppβ2V2R/β-arrestin-1/Fab30 complex formation was disrupted by incubation with excess V2Rpp in a size-exclusion assay. Black: ISO/ppβ2V2R/β-arrestin-1-H295TMSiPhe/Fab30. Red: ISO/ppβ2V2R/β-arrestin-1-H295TMSiPhe/Fab30 complex mixed with 200 μM V2Rpp. The red peak can be simulated by two components, which are as follows: the blue curve represents the SEC of ppβ2V2R alone, and the pink curve represents the V2Rpp/β-arrestin-1-H295TMSiPhe complex. **(f)** 1D ^1^H NMR spectra of β-arrestin-1 H295TMSiPhe in the presence of V2Rpp. The incubation of the V2Rpp with the ISO/ppβ2V2R/β-arrestin-1-H295TMSiPhe/Fab30 complex caused no significant change in the S3 state, but caused a shift of the S1 state to the S2 state. c or f) The data collection buffer used for the experiments contained 20 mM HEPES, 150 mM NaCl, 0.01% LMNG, 0.002 CHS, and 10% D2O (pH 7.5 at 25°C).

As both the receptor core and the phosphorylated receptor C-tail contributed to the interaction between ppβ2V2R and arrestin, we next performed a V2R-phospho-C-tail competition experiment (Figure 5d). Incubation of the excess V2Rpp led to the dissociation of ppβ2V2R from the ISO/ppβ2V2R/β-arrestin-1/Fab30 complex, as suggested by the SEC results (Figure 5e). However, the active conformation of H295 persisted even when the arrestin dissociated from the receptor, as indicated by the maintenance of the amplitude of the S3 state in the 1D ^1^H-NMR spectrum (Figure 5f). These data suggested that the S3 conformational state of H295 in arrestin do not simply reflect the propensities of the ligands for stabilization of the GPCR-arrestin complex, but is a consequence of the receptor-core-arrestin interaction. Furthermore, maintenance of the S3 activation state of arrestin even after dissociation from the receptor was consistent with the hypothesis proposed in recent cellular studies, which suggested that an arrestin activation cycle occurred in response to activation by the receptor.

### Multiple conformational states observed at the ERK interaction site of β-arrestin-1

We next extended the TMSiPhe technology to study the conformations of other β-arrestin-1 sites associated with specific arrestin functions. We selected the R285 position of β-arrestin-1, which was hypothesized to play important roles in interaction with ERK^42^(Figure 3a and Table S3). Notably, superimposition of the structures of inactive β-arrestin-1 structure and the rhodopsin-visual arrestin complex indicated that R285 assumed a highly exposed and extended conformation (Figure 6a), suggesting that receptor interaction may regulate conformational change at this specific site.

**Figure. 6.**
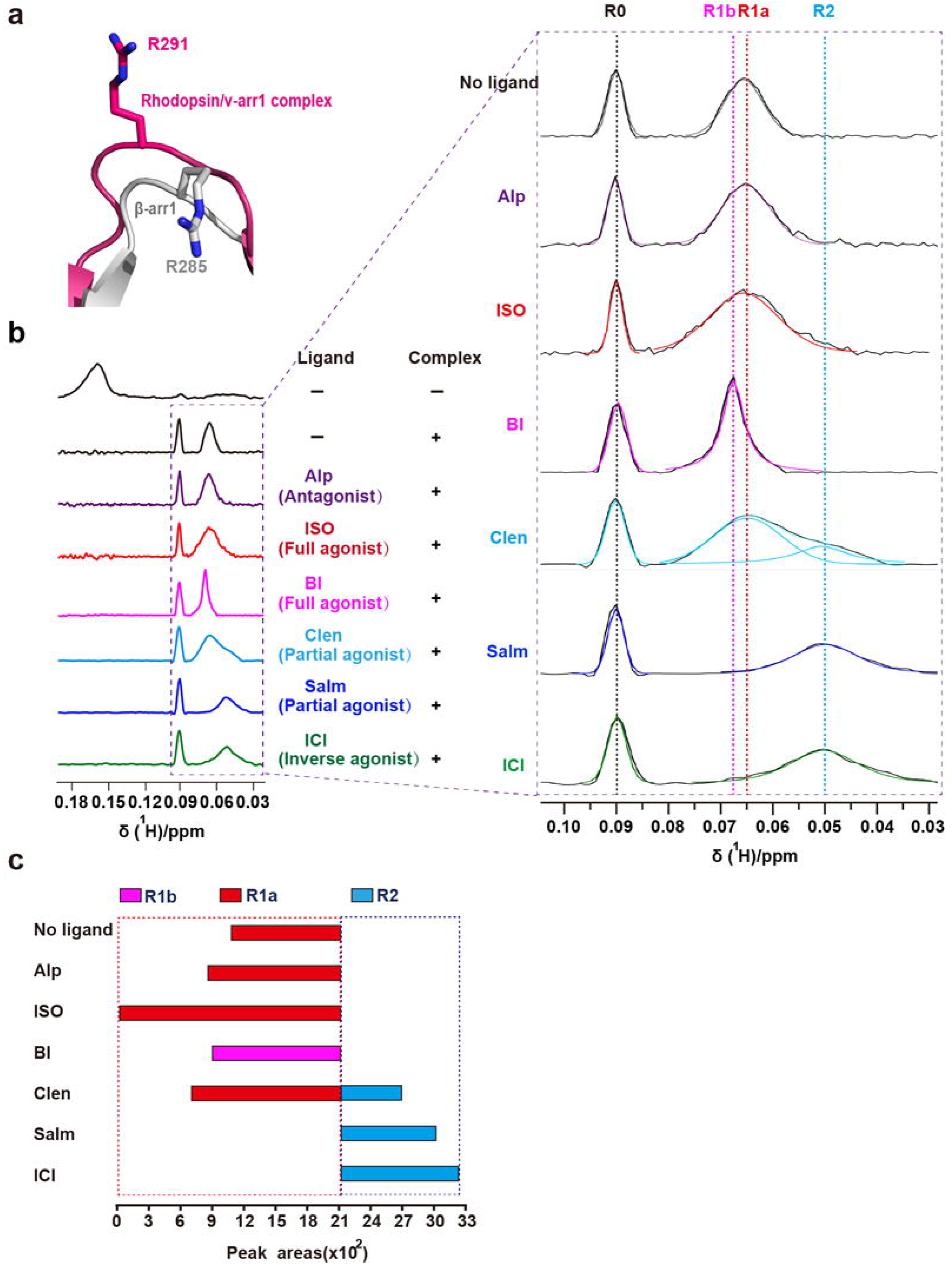
Monitoring the conformational states of site 285 of β-arrestin-1 in response to different β2AR ligands. **(a)**. Structural comparison of the R285 position in inactive β-arrestin-1 (PDB: 1G4M) and the corresponding R291 position in the rhodopsin/arrestin complex (PDB: 5W0P). The active β-arrestin-1 structure is depicted in gray, and the rhodopsin/arrestin complex is in red. The activation of arrestin by a receptor led to a highly solvent-exposed configuration at the R285 position of β-arrestin-1, as suggested by the crystal structures. **(b)** 1D ^1^H NMR spectra of β-arrestin-1 R285TMSiPhe activated by ppβ2V2R with or without different ligands. After incubation with ppβ2V2R, multiple new NMR signals appeared between 0.04 ppm to 0.10 ppm, which are designated as R0 (0.09ppm), R1a (0.065ppm), R1b (0.068 ppm), R2 (0.05 ppm), from low field to high field. The buffer used for the experiment contained 20 mM HEPES, 150 mM NaCl, 0.01% LMNG, 0.002 CHS, and 10% D2O (pH 7.5 at 25°C). **(c)** Bar graph representing the population (simulated peak area) of each NMR peak for each ligand condition. The values are also tabulated in fig. S21.

We therefore incorporated TMSiPhe at the R285 site of β-arrestin-1 and monitored the change in the 1H-NMR spectrum in response to the binding of ppβ2V2R engaged with different ligands (Figure 6b and Figure S19-S21). The functional integrity of R285TMSiPhe- β-arrestin-1 was validated, and Fab30 was used to stabilize the ppβ2V2R/β-arrestin-1 complex without perturbation in the NMR spectrum (Figure S11 and S19). Application of ppβ2V2R without or with different ligands eliminated the original NMR peak at 0.158 ppm but broadened the conformational distributions from 0.03 ppm to 0.10 ppm (Figure 6b-6c and Figure S21). At least 4 different conformational states of the ppβ2V2R/β-arrestin-1-R285TMSiPhe/Fab30 complex were discerned in the presence of different ligands. Notably, β-arrestin-1 alone also has small but visible peaks in the 0.03 ppm-0.10 ppm region, indicating that a conformational selection model may also be suitable for description of the receptor-induced conformational change at the β-arrestin-1-R285 position. In particular, addition of any receptor complexes without or with different ligands all produced a similar peak at 0.09 ppm (R0 state), indicating that this conformational state may be mainly due to the binding of the receptor-phospho-tail but is not significantly affected by the receptor core interaction (Figure 6b and Figure S21).

The ligands mostly changed the distribution of NMR peaks from 0.04 ppm to 0.07 ppm, which included 3 conformational states derived by simulation, namely, R1a-b (0.065-0.068 ppm) and R2 (0.05 ppm). Although application of the neutral antagonist Alp and the agonist ISO had no significant effect on the NMR peak at R1a (0.065 ppm), application of the long-term covalent agonist BI caused a small but significant low field shift of R1a to R1b (0.068 ppm).

The application of partial and inverse agonists caused complex conformational changes. Whereas Clen significantly diminished the distribution of the R1a state and promoted the appearance of a high-field R2 state, the engagement of the receptor with the G-protein-biased partial agonist Sal and the inverse agonist ICI almost completely eliminated the presence of the R1 states and facilitated the emergence of the R2 states (Figure 6b-6c and Figure S21). As Sal and ICI are not known for arrestin-dependent ERK signaling, the appearance of the R2 conformational states of the β-arrestin-1 R285 position may not contribute to ERK activation in response to receptor/arrestin complex interactions. Taken together, multiple conformational states of the β-arrestin-1 R285 position were detected by TMSiPhe in response to different β2AR ligands, which was not strictly correlated with the ability of these ligands in either the activation of G-protein (agonists vs. antagonists) or arrestin-mediated receptor internalization, indicating that each specific receptor ligand may lead to a distinct conformational state at a specific arrestin site, which contributes to the selective functions of these ligands.

## Discussion

Despite the broad applications of NMR in the characterization of protein structure and dynamics, it has remained very challenging to use NMR to study large transmembrane protein complexes, whose NMR spectra exhibit severe line broadening and overlapping resonance. While site-specific protein labelling with 1D NMR probe has provided an exciting new method for the investigation of membrane protein complex^8,9,43–45^, one limitation of cysteine-mediated chemical labeling is that it only allows access to the surface residues of proteins, preventing observation of the important dynamic interactions that occur within protein hydrophobic cores. Moreover, to achieve site-specific labelling, all other surface-exposed cysteine residues must be mutated, which may cause significant perturbation to protein structure and function. By contrast, UAA incorporation through genetic code expansion allows labeling of desired residues at both exposed and internal sites. For example, through genetic code expansion, we have developed a method to efficiently incorporate the UAA difluorotyrosine (F2Y) into proteins of interest, enabling us to study how different receptor phospho-barcodes localized in the receptor C-tail regulate distinct functionally selective arrestin conformations^12,16^. Despite this significant progress, ^19^F-NMR requires large amount of protein (usually more than 100 μM), and each measurement generally takes more than 12 hours. Therefore, the development of a 1D NMR probe for the examination of the conformational dynamics of transmembrane protein complexes using a low concentration of protein is urgently needed.

Through genetic code expansion in *E. coli*, we have achieved the highly selective and efficient labelling of trimethylsilyl (TMS) group in proteins, and demonstrated its broad applicability to investigate multiple conformation state of large membrane protein complexes. The efficient and selective incorporation of TMSiPhe was verified by both mass spectrometry and crystallography. Using this method, we were able to detect the dynamic conformational changes in membrane protein complex (molecular weight ~ 150 kDa) at the residue level, using a low protein concentration ranging of 5 μM, and a short spectra accumulation time of 20 min. Key to this advance is the evolution of TMSiPheRS, a specific tRNA synthetase which selectively recognizes TMSiPhe to facilitate its genetic incorporation into proteins.

Using this method, we were able to observe the ligand-dependent conformational changes in arrestin via direct receptor core engagement, a process important for GPCR signaling. Previous studies by us and others have provided important mechanistic insights, demonstrating that receptor-phospho-barcodes present in the receptor-C-tail play pivotal roles in the determination of selective arrestin functions^12,16–18,32,33,35,46^. An important model for the development of arrestin-biased GPCR ligands is that ligands for GPCRs can cause conformational changes in arrestin via direct receptor core/arrestin interactions regardless of the C-terminal phosphorylation pattern. Notably, the rhodopsin/visual arrestin complex crystal structure provided knowledge of receptor core/arrestin interactions at the atomic level^3^, and the FlAsH-BRET assays revealed that different receptor activation resulted in diverse arrestin conformations in cells^17^. However, dynamic information and high-resolution data regarding conformational changes in arrestin dictated by different receptor ligands via the receptor core/arrestin interaction remains undetermined, likely due to the low resolution of cellular methods and the difficulty of the application of biophysical approaches for the study of receptor complex systems. Here, through the residue-specific conformational detection method using TMSiPheRS, as well as cellular internalization assays, our results reveal that ligands directed structural alterations of the 7-helix transmembrane core of the GPCR interacts with arrestin to cause conformational change in the arrestin polar core, and the extent of which is correlated with the internalization ability of the receptor/arrestin complex.

In addition to structural alterations in the polar core, we used TMSiPhe to examine the conformational changes that occurred at the R285 position of β-arrestin-1, a site associated with ERK activation^42^. Importantly, R285TMSiPhe assumed multiple conformations in response to the engagement of different ligands with β2V2R harboring the same phosphorylated receptor C-tail. Importantly, the conformational states of the 285 site are not directly correlated to the functions of these β2AR ligands in either Gs activation or receptor internalization, indicating that different ligands of the same receptor were able to regulate distinct arrestin conformations at specific arrestin sites, which may be correlated with selective functions. Notably, the arrestin-ERK interaction may involve multiple interfaces. Therefore, the conformational changes in the R285 site observed by TMSiPhe likely contribute to, but are not the sole determinants of, arrestin-mediated ERK activation.

In summary, we have achieved the efficient and selection incorporation of TMSiPhe into protein in *E. coli*, to facilitate rapid detection of the dynamic conformational changes in 150 kD membrane protein complexes, using 1D ^1^H-NMR. Due to the high ^1^H-NMR signal intensity, and unique up-field chemical shift of the TMS group, good 1D ^1^H-NMR spectra can be acquired using only 5 μM of protein, and 20 min accumulation time. Using this handy and powerful approach, we identified the ligand-induced and functionally relevant arrestin conformational states via receptor core engagement^47^. We expect this method will be broadly applicable to biochemistry laboratories to decipher dynamic protein interaction mechanism under physiological conditions.

## Methods

### Reagents

The monoclonal anti-GST (2622), anti-His (2366S) antibodies were purchased from Cell Signaling. The monoclonal anti-Flag M2 antibody (F3165) were purchased from Sigma. Anti -BV envelope gp64 PE antibody (12-6991-80) was purchased from eBioscience. Glutathione-Sepharose 4B and Ni-NTA Agarose were from Amersham Pharmacia Biotech., Isoproterenol, Alprenolol, Clenbuterol, Salmaterol and ICI-118551 were purchased from MCE. BI-167107 was synthesized by Prof. Xin Chen at Changzhou University. V2Rpp were synthesized by Tufts University core facility. Flag M1 antibody were produced by Flag-M1 hybridoma cell and purified by Protein A/G beads. All of the other reagents were from Sigma.

### Constructs

The full-length wild-type cDNAs of bovine β-arrestin-1 was subcloned into the NdeI/XhoI sites of the pET22b vector with the C-terminal His tag. The β-arrestin-1 mutations Y21TAG, Y63TAG, Y173TAG, Y249TAG, R285TAG, H295TAG, L388TAG, sfGFP Y182 TAG were generated using the Quikchange mutagenesis kit (Stratagene). The pFast-β2V2R construct was created by in-fusion of the last 29 amino acid cDNA of human V2-Vasopressin receptor (V2R) into the pFast-β2AR construct has been described previously^12,48^.The pcDNA3.1-Flag-β2V2R-Rluc was created by in-fusion of the Rluc plasmid with the pcDNA3.1- Flag-β2V2R construct. All constructs and mutations were verified by DNA sequencing.

### Synthesis of TMSiPhe

The synthesis of TMSiPhe according to the route in fig. S2, with following steps^26^.

Synthesis of trimethyl(4-tolyl) silane (2).

Iodine (catalytic amount) was added to the mixture of Magnesium turning (2.67 g, 110 mmol) and 4-bromotoluene 1 (1.71 g, 10 mmol) in 80 ml of dry tetrahydrofuran (THF (containing 0.002% water). The reaction was started by heating, then 4-bromotoluene (1) (15.4 g, 90 mmol, dissolved in 20 mL of dry THF) was slowly added in a drop wised manner. After refluxing for 4h, the reactions were kept slight boiling by the drop wised addition of trimethyl chlorosilane (12.7 ml, 110 mmol). The mixture were reflux for another 2 h, followed by stirring at room temperature and quenching with 500 ml ice-cold water. The mixture was extracted with ethyl acetate (EA, 100 mL*3) and the organic layers were combined and subsequently washed with brine (100 mL*3). The organic layer was then dried over Na2SO4, filtered and evaporated. The residue was chromatographed by silica gel with petroleum ether (PE) as an eluent. The colorless liquid (14.3 g) was obtained with 87% yield.

1H NMR (500 MHz, CDCl3) δ 7.45 (d, J = 7.6 Hz, 2H), 7.21 (d, J = 7.4 Hz, 2H), 2.38 (s, 3H), 0.28 (s, 9H).

Synthesis of (4-(bromomethyl) phenyl) trimethylsilane (3)

Trimethyl(4-tolyl) silane (2) (3.28 g, 20 mmol) was dissolved in tetrachloromethane (CCl4, 50 mL, A.R. grade) at room temperature. N-bromosuccinimide (NBS, 3.56 g, 20 mmol) and azodiisobutyronitrile (AIBN, 0.33 g, 2 mmol) was added. The mixture was stirred with 4hours refluxing, followed by vacuum condensation. The residue was used for the next step without further purification.

1H NMR (500 MHz, CDCl3) δ 7.51 (d, J = 7.9 Hz, 2H), 7.38 (d, J = 7.9 Hz, 2H), 4.51 (s, 2H), 0.28 (s, 9H).

Synthesis of ethyl 2-((diphenylmethylene)amino)-3-(4-(trimethylsilyl) phenyl) propanoate (4)

N-(Diphenylmethylene)glycine ethyl ester (13.37 g, 50 mmol) and potassium hydroxide (8.42 g,150 mmol) was dissolved in 60 ml DMSO and the mixture was stirred at 10℃ for 20 min. The mixture was added with (4-(bromomethyl) phenyl) trimethylsilane (3) (12.15 g, 50 mmol) and kept stirring for 1 h, following by adding 720 ml of ice-cold water and then extracting with EA (200 ml*3). The organic layers were combined and were subsequently washed with brine (100 mL*3). The organic layer was then dried over Na2SO4, filtered, and concentrated under reduced pressure.

Synthesis of ethyl 2-amino-3-(4-(trimethylsilyl) phenyl) propanoate (5)

The residue from preceding step was added with THF 60 ml and 1N HCl aqueous 60ml. The solution was stirred for 1 h and then was added with 180 ml of PE, washed with PE/diethyl ether (3:1) (200 ml*3). The organic phase was extracted with 0.1N HCl aq (100 ml*3). Then the aqueous phase was combined and alkalized with Na2CO3 to pH= 9~10 and extracted with EA (100 mL*3). The final organic layers were combined and subsequently washed with brine (100 mL*3), dried over Na2SO4 and concentrated. 8.1 g compound 5 was acquired finally. The yield for the product is approximately 60% over these 3 steps.

1H NMR (500 MHz, CDCl3) δ 7.44(d, J = 7.7 Hz, 2H), 7.27 (d, J = 7.7 Hz, 2H), 4.43 (s, 1H), 4.14 (q, J = 6.8 Hz,2H), 3.49 (m, 1H), 3.38 (m, 1H), 1.15 (t, J = 6.9 Hz, 3H), 0.24 (s, 9H).

Synthesis of 2-amino-3-(4-(trimethylsilyl) phenyl) propanoic acid (6) 7.9 g of Ethyl 2-amino-3-(4-(trimethylsilyl)phenyl)propanoate (5)(30 mmol)was added with THF 30 ml and 2N NaOH aqueous 30 ml. The mixture was then stirred for overnight at room temperature, followed by adding 300 ml of PE. Then the aqueous phase was added to 600 ml of 0.1N HCl aq in a drop wise manner with stirring. A lot of white solid was precipitated from the solution. The product was filtered and dried under vacuum to afford the 2-amino-3-(4-(trimethylsilyl) phenyl) propanoic acid (5.6 g, 78%).

1H NMR (500 MHz, D2O) δ 7.48 (d, J = 6.6 Hz, 2H), 7.19 (d, J = 6.6Hz, 2H), 3.39 (m, 1H), 2.91 (m,1H), 2.72(m,1H), 0.14 (s, 9H).13C NMR (100 MHz, MeOD-d3) δ172.28, 140.70, 136.82, 135.05, 129.88, 55.99, 37.66, −1.13.

HRMS (ESI) calculated for [M+H]+C12H20NO2Si: 238.1258, found 238.1256.

### Genetic selection of the mutant synthetase specific for TMSiPhe (TMSiPheRS)

The pBK-lib-jw1 library consisting of 2×109 independent TyrRS clones was constructed using standard PCR methods. E. coli DH10B harboring the pREP(2)/YC plasmid was used as the host strain for positive selection. Cells were transformed with the pBK-lib-jw1 library, recovered in SOC for 1 h, washed twice with glycerol minimal media with leucine (GMML) before plating on GMML-agar plates supplemented with kanamycin, chloramphenicol, tetracycline and TMS-Phe at 50 g/ml, 60 g/ml, 15 g/ml and 1mM respectively. Plates were incubated at 37 °C for 60 hours and surviving cells were harvested. Subsequently, the plasmid DNA was extracted and purified by gel electrophoresis. The pBK-lib-jw1 DNA was then transformed into electro-competent cells harboring the negative selection plasmid pLWJ17B3, recovered for 1 h in SOC and then plated on LB-agar plates containing 0.2% arabinose, 50 g/ml ampicillin and 50 g/ml kanamycin. The plates were then incubated at 37 °C for 8-12 hours, and pBK-lib-jw1 DNA from the surviving clones was extracted as described above. The library underwent another round of positive selection, followed by a negative selection and a final round of positive selection (with chloramphenicol at 70 g/mL). At this stage, 96 individual clones were selected and suspended in 50 L of GMML in a 96-well plate, and then replica-spotted on two sets of GMML plates. One set of GMML-agar plates was supplemented with tetracycline (15 g/mL), kanamycin (50 g/mL) and chloramphenicol at concentrations of 60, 80, 100 and 120 g/mL with 1 mM TMSiPhe. The other set of plates were identical but did not contain TMSiPhe, and the chloramphenicol concentrations used were 0, 20, 40 and 60 g/mL. After 60 h incubation at 37 °C, one clone was found to survive at 100 g/mL chloramphenicol in the presence of 1 mM TMSiPhe, but only at 20 g/mL chloramphenicol in the absence TMSiPhe.

### Purification of TMSiPheRS

TMSiPheRS was purified from E.coli as described previously^22^. Briefly, the gene encoding the TMSiPheRS was cloned into the pET22b vector and then transformed into BL21(DE3) cells. The large scale expression cultures were grown to an OD of 0.8. After induction for 4-6 hours at 37℃ with 1 mM IPTG, cells were pelleted by centrifugation and re-suspended in lysis buffer (50 mM Tris, pH 8.5, 500 mM NaCl, 10 mM β-mercaptoethanol, 5 mM imidazole). Cells were sonicated and the cell lysate was pelleted by centrifugation. The supernatant was collected and incubated with Ni-NTA agarose beads for 2 hours at 4°C, filtered, and washed with wash buffer (50 mM Tris, pH 8.5, 500 mM NaCl, 10 mM β-mercaptoethanol, 20 mM imidazole). The synthetase was eluted with a wash buffer containing 300 mM imidazole in buffer A (25 mM Tris, pH 8.5, 25 mM NaCl, 10 mM β-mercaptoethanol, 1 mM EDTA), purified by anion exchange chromatography (Hitrap MonoQ; GE Healthcare) using a salt gradient from 25 mM to 0.5 M NaCl. TMSiPheRS was purified by Sephadex gel column chromatography (Superdex 200 10/300 GL; GE Healthcare) in a buffer containing 50 mM Tris, pH 8.5, 500 mM NaCl, 10 mM β-mercaptoethanol and concentrated to 25 mg/mL.

### Preparation crystals for TMSiPhe incorporated sfGFP

The plasmids encoding sfGFP Y182TMSiPhe in pET22b vector was co-transformed with pEVOL-TMSiPheRS into BL21(DE3) E.coli cells. Cells were amplified in LB media supplemented with ampicillin (50 μg/mL) and chloramphenicol (30 μg/mL). Cells were then grown to an OD600 = 0.8 at 37°C. After induction14 hours at 30°C with 0.2% L-arabinose, 0.3 mM IPTG and 0.5 mM TMSiPhe, cells were harvested by centrifugation. The cells were lysed by French pressing in buffer containing 50 mM HEPES, pH 7.5, 500 mM NaCl. The supernatant was collected and incubated with Ni-NTA column for 2 hours at 4°C, filtered, and washed with wash buffer containing 50 mM HEPES, pH 7.5, 500 mM NaCl, 20 mM imidazole. The protein was eluted with a wash buffer containing 50 mM HEPES, pH 7.5, 500 mM NaCl, 250 mM imidazole. sfGFP Y182TMSiPhe was purified by size exclusion column (Superdex 200 increase 10/300 GL; GE Healthcare) in a buffer containing 20 mM HEPES-Na, pH 7.5, and concentrated to 20 mg/mL. The crystal of sfGFP Y182TMSiPhe were obtained at 16℃ by the hanging drop vapor diffusion by mixing 1 μL protein sample with equal volume of mother liquor containing 10% PEG 6,000 and 2.0 M Sodium chloride. The crystal appeared within one week. Crystals were then flash-frozen in liquid nitrogen in 10% PEG 6000, 2.0 M Sodium chloride and 20% glycerol.

### Data collection and Structure determination of TMSiPhe incorporated sfGFP

Diffraction data for sfGFP Y182TMSiPhe were collected at beamline BL19U1 of Shanghai Synchrotron Radiation Facility (SSRF). All data collected were indexed, integrated and scaled using software of XDS and Aimless respectively^49,50^. The structure of sfGFP Y182TMSiPhe was solved by molecular replacement using sfGFP-66-HqAla, (PDB code: 4JFG) as a search model by Phaser within PHENIX package. Structural refinement was carried out by Phenix. In the refinement process, the program Coot in the CCP4 program suite was used for the model adjustment, and water finding, whereas ligand restraints were produced using the eBLOW contained in PHENIX software package^51^. The structure models were checked using the PROCHECK^56^.

### Preparation crystals for TMSiPheRS alone and TMSiPheRS complex

Crystals of TMSiPheRS alone were grown at 16°C using the hanging drop vapor diffusion technique against a mother liquor composed of 22% polyethylene glycol (PEG) 1500, 100 mM Hepes (pH 7.5) and 200 mM L-Proline and 1:1 mixture of concentrated synthetase (25 mg/mL). For TMSiPheRS complex, TMSiPhe (100 μM) was incubated with TMSiPheRS (10 μM) for 2 hours at 25°C. The complex was concentrated to 20 mg/ml, Crystals were grown in hanging drops containing 1.5 μL of complex solution and 1.5 μL of a well solution composed of 24% PEG1500, 100 mM Hepes (pH 7.5) and 200 mM L-Proline. The crystal appeared after about one week. Crystals were flash frozen in liquid nitrogen after a 30s soak in 26% PEG 1500, 100 mM Hepes (pH 7.5) and 200 mM L-Proline and 20% glycerol.

### Data collection and Structure determination of TMSiPheRS alone, TMSiPheRS complex

X-ray diffraction data of TMSiPheRS alone and TMSiPheRS complex were collected at beamline BL19U1 of Shanghai Synchrotron Radiation Facility (SSRF). All data collected were indexed, integrated and scaled using software of XDS and Aimless respectively^49,50^. The structure of TMSiPheRS alone and TMSiPheRS complex was solved by molecular replacement using F2Y–F2YRS complex(PDB code: 4HJX) as a search model by Phaser within PHENIX package. Structural refinement was carried out by Phenix. In the refinement process, the program Coot in the CCP4 program suite was used for the model adjustment, and water finding, whereas ligand restraints were produced using the eBLOW contained in PHENIX software package^51^. The structure models were checked using the PROCHECK^56^.

### Peptide synthesis

A fully phosphorylated 29-amino-acid carboxy-terminal peptide derived from the human V2 vasopressin receptor (V2Rpp: 343ARGRpTPPpSLGPQDEpSCpTpTApSpSpSLAKDTSS371) was synthesized from Tufts University Core Facility. And the high-affinity version of Gtα(340ILENLKDCGLF350, GtαCT-HA) were purchased from China Peptides Co., Ltd. with more than 95% purity as verified by analytical high-performance liquid chromatography. In the competition assays, the GtαCT and the V2Rpp were used as 200 μM concentration.

### Expression and purification of β-arrestin-1 TMSiPhe mutants

The pEVOL-TMSiPheRS plasmids encoding specific M. jannaschii tyrosyl amber suppressor tRNA/tyrosyl-tRNA synthtase mutants were co-transformed into E. coli BL21 (DE3) together with the pET22b vector harboring the target β-arrestin-1 mutant. The E. coli cells were cultured in Luria-Bertani (LB) medium. After the 1L cell culture reached OD600 0.6-0.8, the cells were induced with 300 μM isopropyl-β-D-thiogalactopyranoside (IPTG) and 0.2% L-arabinose for 12 h (25℃) to allow protein expression in presence of 0.5mM TMSiPhe in the culture medium. The cells were lysed by French pressing in buffer A (50 mM Tris-HCl, pH 8.0, 150 mM NaCl) and the lysate was batch binding with 300 μL Ni-NTA column (GE Healthcare, USA). After an extensive washing with buffer A, the target protein was eluted using 300mM imidazole in buffer A. These proteins were subsequently purified by size exclusion column Superdex 75 and the buffer was exchanged to buffer B (50 mM Tris-HCl, pH 7.5, 150 mM NaCl).

### Expression and purification of β2V2R

FLAG-β2V2R and GRK2-CAAX were co-expressed in baculovirus-infected insect cells (Sf9) using the Bac-to-Bac baculovirus Expression System as previously described^38^. Cells were stimulated with ISO(10 μM)and harvested at 64 or 72 h after infection. The cell pellets were stored at –80℃. Cell membranes were disrupted by thawing frozen cell pellets in 300 ml of hypotonic buffer C (10 mM HEPES pH7.5, 20 mM KCl and protease inhibitor cocktail) and homogenized using a Dounce homogenizer repeated plunging. The membrane fraction was separated from the lysate via ultracentrifugation (42,000 rpm speed for 40 min in Ti45 rotor). The pellet was washed 3-4 times with a high osmotic buffer D containing 1.0 M NaCl in the above buffer C, and centrifuge as above. The pellet was subsequently solubilized with 1% n-decyl-β-D-maltopyranoside and (DDM, Anatrace) 0.2%CHS (sigma) in buffer E (50 mM HEPES pH7.5, 1 M NaCl). The solubilized membrane fraction was then purified by flag-M1 resin (sigma) affinity chromatography in buffer F (20 mM HEPES pH7.5, 150 mM NaCl, 0.1%DDM, 0.02% CHS). Finally, the sample buffer was exchanged to buffer G (20 mM HEPES pH7.5, 150 mM NaCl, 0.01%LMNG, 0.002% CHS) using a PD-10 desalting column. Purified protein samples were used fresh in the experiments.

### Superdex Exclusion Chromatography

The purified ppβ2V2R (30 μM) were stimulated with different ligands (60 μM) and then incubated with β-arrestin-1 H295TMSiPhe (10 μM) for 30 min at 25°C. Then Fab30 (20 μM) was then added to the mixture and the complex was allowed to form for 1h at 25 °C. The ligand/ppβ2V2R/β-arrestin-1 H295TMSiPhe-Fab30 complex were concentrated and then purified by Superdex 200 increase in 20 mM HEPES pH7.5, 150 mM NaCl, 0.01% LMNG, 0.002% CHS and corresponding ligand (60 μM). The yield of the purified complexes were approximately 50%, and the purities were judged by size exclusion chromatography and the electrophoresis.

### NMR experiment

β-arrestin-1 TMSiPhe mutants prepared for NMR analysis were quantified with BCA protein assay kit and diluted with buffer B (containing10% D2O)to 5~20 μM. All 1D 1H NMR spectra were recorded with typical total experimental times 8~15 min at 25°C, on an Avance 950 MHz spectrometer with cytoprobe (Bruker, Billerica, MA). The spectra were processed and analyzed with the program ZGGPW5 (NS = 32; DS = 4; SW = 20ppm; AQ = 1.93 s; D1 = 1s. The number of scans was adjusted to the relative protein concentration in each experiment. The chemical shift of the signal peak was determined by reference to D2O (4.68ppm).

Binding of the V2Rpp to the β-arrestin1 was assessed using β-arrestin1-F388TMSiPhe (20 μM), in the presence of V2Rpp at a gradient increased concentration, in 50 mM Tris-HCl, pH 7.5, 150 mM NaCl, 10% D2O buffer on a Bruker 950 MHz NMR spectrometer. The signal was normalized with Tris and integrated at shift −0.05 ppm after auto baseline correction by MestReNova.9. Through calculating the ratio of the area of remaining Apo NMR peak and the original concentration of each component, the complex state (Bound) concentration and free ligand(V2Rpp)concentration were obtained for Scatchard plotting and one-site specific curve fitting.

Buffer for complex of ppβ2V2R/β-arrestin1/Fab30 1D1H NMR spectra was 20 mM HEPES, 150 mM NaCl, 0.01% LMNG, 0.002% CHS, 10% D2O, pH 7.5, 60 μM ligand or control vehicle (diluted DMSO). The total recording time for each experiment was 40 min. spectra were recorded using a Bruker 950 MHz NMR spectrometer at 25°C.

### Expression and purification of Fab 30 proteins

The purification of Fab30 was performed as previously described^36^. M5532 E. Coli competent cells was transformed with the plasmid containing Fab30 fragment and was cultured in the CRAP-Amp medium cultures in 2.8 L non-baffled flasks and grow for 18-24 hours at 30°C (200 rpm). These cells were pelleted and freeze with liquid nitrogen, then stored at −80°C. The frozen cell pellets were thawed at room temperature and added with 15 mL of TES (Tris-EDTA-Sucrose) / pellet of 1 liter culture; resuspend, shaked for one hour on ice in cold room platform shaker with 150 rpm (TES buffer: 200mM Tris pH=8.0; 0.5 mM EDTA, 0.5 M sucrose), followed by adding with 30 ml of TES/4 (TES one part plus 3 parts ice cold deionized H2O) per pellet of 1 liter culture and continue to shake 1h on ice. The solution was poured into 250 mL centrifuge bottles and spin in SLA 1500 rotor for 30 minutes at 15000 rpm. All remaining purification steps were carried out in cold room. The supernatant of the cell lysate were incubated with Ni-NTA beads by 2-12 hours with a ratio of 500 μL beads/ 1 liter culture. The beads were packed in a column and washed with 40 CV of cold buffer B (20 mM Tris-Hcl pH=7.55, 150 mM NaCl), and then eluted with buffer C (20 mM Tris-HCl pH=7.55, 150 mM NaCl, 250 mM imidazole).

### GST pull down assay

0.1 μM wild-type or mutant β-arrestin-1 was mixed with 0.5μM phospho-receptor-C-tail fragment (V2Rpp) and incubated in binding buffer (20 mM Tris-HCl, pH 7.5, 150 mM NaCl, 2 mM EDTA, 1 mM DTT) at 25°C for 30 min as previously described^12^. 1 μM GST-clathrin was then added and incubated for another hour. Subsequently, 10 μL GST beads were added into the mixture and the mixture was rolled at 4°C for 2 h. The GST beads were collected by centrifuge and washed with wash buffer (binding buffer with 0.5% Tween20) for 4 times. After removing the supernatant, the samples were re-suspended in 50 μL 2×SDS loading buffer and boiled for 10 min before western blot.

### Bimane labeling of purified receptors and TRIQ experiment

The method was carried out according to a previously published manuscript^52^. Purified receptors (β2AR-Δ5-Cys271+Trp135)and mBBr (Invitrogen) were mixed at the same molarity in LMNG buffer (not containing CHS) and incubated overnight on ice in the dark. Fluorophore-labeled receptors were obtained by gel filtration on a desalting column equilibrated with LMNG/CHS buffer (20 mM HEPES, 150 mM NaCl, 0.01% LMNG, 0.002% CHS, pH 7.5).

Fluorescence spectroscopy was measured on a Varioskan flash (Thermo Scientific) instrument with full wavelength scanning mode at 25°C. 100 μL samples containing 0.2 μM bimane labeled β2AR in a MicroFluor 96-well plate were excited at 390 nm, and the emission fluorescence was measured by scanning from 430 to 500 nm using a 2 nm step. Each data point was integrated for 0.2 s. If the ligand was present, concentration of ligand was set for 5μM and the incubation time was set for 15min.We corrected fluorescence intensity for background fluorescence from buffer. Spectra were analyzed using the GraphPad Prism 5.

### BRET assay

HEK293 cells seeded in 6-well plates were transfected with 0.5 μg BRET donor Flag-β2V2R-Rluc and 1 μg BRET acceptor Lyn-YFP using polyethylenimine as previously described^21^. 24 h after transfection, the cells were detached and distributed into 96-well plates at a density of ~25,000 cells per well. After another 24 h incubation at 37 °C, the cells were washed twice with Tyrode’s buffer (140 mM NaCl, 2.7 mM KCl, 1 mM CaCl2, 12 mM NaHCO3, 5.6 mM D-glucose, 0.5 mM MgCl2, 0.37 mM NaH2PO4 and 25 mM HEPES, pH 7.4) and stimulated with vehicle or different ligands (final concentration of 10 μM) at 37 °C for 20 min. Luciferase substrate coelenterazine-h was added at a final concentration of 5 μM before light emissions were recorded using a Mithras LB940 microplate reader (Berthold Technologies) equipped with BRET filter sets. The BRET signal was determined by calculating the ratio of the light intensity emitted by YFP (530/20 nM) over the light intensity emitted by Rluc (485/20 nM).

### Intra-gel digestion and LC-MS/MS analysis and database search

The TMSiPhe incorporated β-arrestin1 was purified and subjected to the electrophoresis. After decolorized, DTT reduction and alkylated by iodoacetamide, the dyeing strip was digested by trypsin overnight. The peptides were extracted with 60% acetonitrile. The peptide mixture obtained after enzymatic hydrolysis was analyzed by a liquid chromatography-linear ion trap-orbitrap (nanoLC-LTQ-Orbitrap XL, Thermo, San Jose, CA) mass spectrometer. The chromatographic column was a C18 reverse phase column. Mobile phase A: 0.1% FA/H2O, B: 0.1% FA/80%CAN/20% H2O, flow rate 300 nL/min. A gradient of 90 min was used. Data analysis was performed using Proteome Discoverer (version 1.4.0.288, Thermo Fischer Scientific) software. The MS2 spectrum uses the SEQUEST search engine to search for arrestin H295TMSiPhe containing fasta. The search parameters are: Trypsin enzymatic hydrolysis, half cut, two missed cut sites, precursor ion mass error less than 20 ppm, and fragment ion mass error less than 0.6 Da. The alkylation of cysteine was set as a fixed modification, and the oxidation of methionine and the specific modification of histidine (H+82.049 Da) were variable modifications. The retrieved peptides and spectral matches (PSM) were filtered using the Percolator algorithm with a q value of less than 1% (1% FDR). The retrieved peptides are combined into a protein under strict maximum parsimony principles.

### Q-TOF mass spectrometry spectrum analysis and database search

LC-MS analysis was performed using a Agilent Q-TOF mass spectrometer in line with a Agilent 1290 HPLC system. The 5 μl purified TMSiPhe incorporated β-arrestin1 protein was loaded onto a reverse phase column (30 0SB-C8, 2.1 x 50 mm, 3.5 μM particle) (Agilent Technologies, SantaClara, CA). The proteins were then eluted over a gradient: 2% B for 2min to waste, then turned LC to MS, 2–50% B in 6 min, 50–90% B in 4 min, 90% B sustained for 4min, then decreased to 2% in 1.1 min, (where B is 100% Acetonitrile, 0.1% formic acid, A is water with 0.1% formic acid) at a flow rate of 0.2 mL/min.and the elution was introduced online into the Q-TOF mass spectrometer (Agilent Technologies, SantaClara, CA) using electrospray ionization. MS data were analysed by MassHunter biocomfirm software.

### Statistical Analysis

For all experiment, the number of replicates and P value cutoff are described in the respective figure legends. Error bars are shown for all data points with replicates as a measure of variation with the group. Statistical differences were determined by One-way ANOVA using the analysis software GraphPad Prism (*P<0.05; **P<0.01;***P<0.005)

## Acknowledgements

NMR experiments were performed at the performed at the Beijing NMR Center and the NMR facility of National Center for Protein Sciences at Peking University, and Nuclear Magnetic Resonance Science Research Platform at Wuhan Institute of Physics and Mathematics, Chinese Academy of Sciences (CAS). We thank the staff of BL19U1 beamlines at National Center for Protein Sciences Shanghai and Shanghai Synchrotron Radiation Factility for assistance during data collection. We thank Prof. Mai-li Liu and Prof. Xu zhang from Wuhan Institute of Physics and Mathematics, CAS, Hong-wei Li from Beijing nuclear magnetic resonance center, Ms Shan-shan Zang and Dr Xue-hui Liu from the Core Facility of Protein Research, Institute of Biophysics (IBP), CAS, for their help in the NMR data collection, analysis and valuable discussion. We thank Jun-ying Jia and Shuang Sun from Lab of Structural and Functional Analysis (IBP, CAS) for their technical assistance in flow cytometry analysis, and their colleague Jian-Hui Li for assistance in fluorescence measurement. We thank Xiang Ding and Zhen-sheng Xie from Lab of Proteomics (IBP, CAS) for LC-MS analysis.

We acknowledge support from the National Key Basic Research Program of China (2018YFC1003600 to J.-P.S; 2017YFA0505400, 2016YFA0501502 to J.Y.W. and F. H. L.), the National Natural Science Foundation of China (81773704 and 31611540337 to J.-P.S., 21837005, 21750003 to J.Y.W., U1532150 and 91640106 to F. H. L., 31700692 to P.X.), The National Science Fund for Distinguished Young Scholars (81825022 to J.-P.S.) and the Program for Changjiang Scholars and Innovative Research Team in University (IRT13028).

## Author Contributions

J.-P.S. conceives the idea for extracellular ligands induced conformational changes in β-arrestin-1 via direct receptor 7-transmembrane core interactions. J.Y.W. conceived the idea that TMSiPhe should be an excellent nuclear magnetic probe, and designed evolutionary strategy for TMSiPheRS. J.-P.S., J.Y.W. and X.Y. designed most of the experiments. Q.L., F.Y., P.S., Q.-W.W. and F. Z. collected and analyzed the 1H NMR data. X.-X.L. and F.-H.L. realized insertion and verification of unnatural amino acids. Z.-L. Z. and Q.-T.H. Performed crystallization and structure solution and analysis of the synthetase and TMSiPhe decorated GFP. Q.L., X.-Y.W., F.Y., P. X., S.-M.H., S.-C.G. and M.-J.H. expressed and purified GPCR proteins. Q.-T.H. carried out Superdex Exclusion Chromatography. Q.-T. H.and C.-X.Q. supplied critical antibody Fab 30. Q.L., Z. X., and S.-L.S. Synthesized TMSiPhe. Prof. X.C. synthesized the compound BI-167107. Z. Y. performed BRET assay. Z.-Y.Y. performed GST pulldown assay. Q.L. performed fluorescence spectroscopy assay. Q.L., Z. G. and J.-Y. L. performed FSEC analysis for GPCR-arrestin complex (Data not shown). A.W. K., K.-H.X. and J.-P.S. decorated the GFP on the arrestin and developed the FSEC methods. K. R., X.-G.N. and C.-W. J. participated in the design and explanation of the NMR results. W. K., X.Y. supervised the fluorescence and BRET experiments.

J.-P.S., J.Y.W. and X.Y. supervised the overall project design and execution. J.-P.S. participated in data analysis and interpretation. Professor K.K.Z. and K.R. offered important advice and help especially in NMR experiment. J.-P.S. and J.Y. W wrote the manuscript. All of the authors have seen and commented on the manuscript.

